# Stem bending generates electrical response in poplar

**DOI:** 10.1101/2020.10.19.345959

**Authors:** Erwan Tinturier, Éric Badel, Nathalie Leblanc-Fournier, Jean-Louis Julien

## Abstract

Under natural conditions, plants experience external mechanical stresses such as wind and touch that impact their growth. A remarkable feature of this mechanically induced growth response is that it may occur at distance from the stimulation site, suggesting the existence of a signal propagating through the plant. In this study, we investigated the electrical response of poplar trees to a transient controlled bending stimulation of the stem that mimics the mechanical effect of wind. Stem bending was found to cause an electrical response that we called ‘gradual’ potential, similar in shape to an action potential. However, this signal distinguished from the well-known plant action potential by its propagation up to 20 cm along the stem and its strong dumping in velocity and amplitude. Two hypotheses regarding the mode of propagation of the ‘gradual’ potential are discussed.

**One sentence summary:** Poplar stem bending induces an electrical response with high speed and strong decrement.

## Introduction

Plants perceive mechanical disturbances from their environment and adjust their growth according to a process called thigmomorphogenesis (Jaffe, 1973). In natural conditions, trees experience wind every day, applying recurrent force loadings that lead to mechanical strain of stem and branches. As a physiological response, trees in windy environments show a low primary growth and an increase of their diameter (Telewski, 2006). The responses to mechanical disturbances can be local or remote. As shown by Coutand et al. (2000), bending of a tomato stem leads to a rapid cessation of primary growth away from the stimulated area. Although, this rapid response has not been confirmed in poplar after a single basal flexion of the stem (Tixier et al., 2014), a slowdown in primary growth was observed following daily repeated flexions for weeks (Niez et al., 2019). These observations suggest the ability of the plants to transport mechanically-induced information over a long distance.

Several long-distance signaling ways can be considered from the literature essentially studied after wounding stimuli: chemical signaling (Choi et al., 2016), hydraulic pulse (Malone, 1993; Lopez et al., 2014) and electrical signaling (Mousavi et al., 2013; Hedrich et al., 2016). Although these pathways are generally studied separately, they are likely to interact (Farmer et al., 2014; Van Bel et al., 2014). The electrical signaling hypothesis has not yet been studied in case of non-wounding mechanical stimulations such as organ bending triggered by wind.

In plants, two types of electrical signals were described: the action potential (AP) and the slow wave (SW) or variation potential. The SW is characteristic of a response to a solicitation producing wounds. It is consisted of a rapid depolarization phase followed by a short plateau step where the transmembrane potential remains stable, and finally a repolarization step lasting variably from 1 to 45 minutes. Its propagation speed is of the order of mm.s^−1^ (Sambeek and Pickard, 1976; Roblin and Bonnemain, 1985; Julien et al., 1991). The SW is positively correlated with stimulus intensity and its amplitude decreases with increasing distance from the stimulation site (Stahlberg et al., 2005). Several studies reported the SW involvement in the activation of defense mechanisms such as the proteinase inhibitor induction (Wildon et al., 1992) and jasmonates signaling pathway (Mousavi et al., 2013). The AP is a rapid and transient depolarization (a few seconds to 2 min) of the plasma membrane with all-or-nothing characteristics, propagating with a tissue-specific rate and without decrement (Zawadzki et al., 1991; Fromm and Spanswick, 1993; Stankovic et al., 1998). The signal propagates preferentially through the phloem, which shares common properties with neurons in animals, namely the existence of membrane continuity by which electrical excitations could be transmitted from one cell to another via plasmodesmata, pores connecting the cytoplasm of one cell to that of its neighbour (Fromm and Lautner, 2007; Van Bel et al., 2014). AP has mainly been studied in the context of rapid motor activity of plants, such as carnivorous or *Mimosa pudica* plants, in which a propagation rate of the order of 5 cm.s^−1^ has been measured (Sibaoka, 1969). APs have also been triggered in plants without rapid motor activity such as *Lupinus* or willow (Paszewski and Zawadzki, 1976; Fromm and Spanswick, 1993).

The rapid propagation of AP could explain the rapid onset of remote responses such as longitudinal growth inhibition observed in plants after stem bending. As highlighted by recent studies, trees are particularly impacted by mechanical stimulations related to wind (Alonso-Serra et al., 2020). Therefore, the aim of this study is to investigate the electrical signals triggered after a transient bending of poplar stems by electrophysiological measurements.

## Results

To test the effect of stem bending on the production of an electrical signal that propagates, the first challenge was to develop an original device (Fig. 1) in which the measuring electrodes were kept immobile during the stem bending. In our case, we attached the stem to two fixed points. This mechanical configuration avoided the possible rotation of the cross sections of the stem and thus, the movement of its non-solicited part. This ensured that the electrodes were kept totally immobile when the stem was bent.

**Figure 1:**
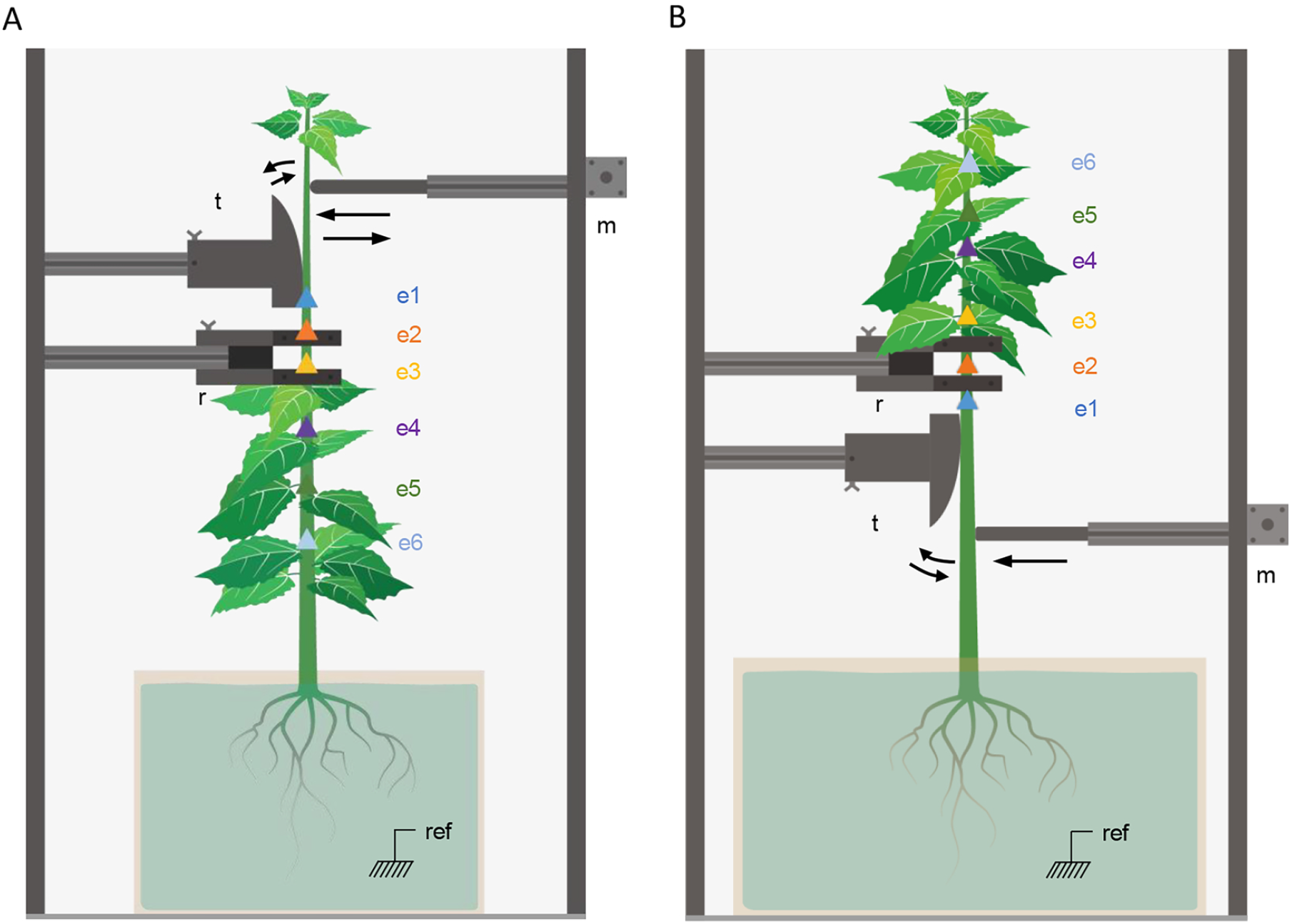
Experimental designs for measuring electrical signals in stem after an apical (A) or a basal (B) transient bending. The poplar tree is placed in a Faraday cage; fxed in two points of the stem with clamping rings (r). The roots incubate in a nutrient solution. The stem is bent at 2.5 cm.s-1 with a motorized arm (m) that pushes the stem along a plastic constantcurvature template (t). The electrodes are placed in the immobile portion of the stem. Electrode e1 is inserted 2 cm below the stem-template contact. Then the distances are: e1-e2 = 2 cm, e2-e3 = 3 cm, e3-e4 = 5 cm, e4-e5 = 5 cm and e5-e6 = 5 cm. The potential difference is monitored along 20 cm of straight stem, below (A) or above (B) the bent region.

A single, transient and rapid bending (2s) of the apical part of poplar stem induced a depolarization wave (Fig. 2A) propagating basipetally up to 20 cm with a significant decrement. The mean amplitude near the flexion point was around 34 mV (e1), and dropped to 24 mV after 5 cm (e3), to reach a mean close to zero at 20 cm (e6 = 2.6 mV) (Fig. 2B). The passage of the depolarization wave was followed by a slight hyperpolarization of the four measuring electrodes closest to the stimulation. Then, a slow (up to 30 minutes) and oscillation-free return to the initial potential was observed. The half-life time of the depolarization wave also decreased with the position of the electrode; from 44.2 s for e1 (the closest electrode from the bent zone) to 34.3 s for e3 and 6.8 s for e6, the farthest electrode (Table 1A). The average propagation speed of the depolarization wave was evaluated between each electrode (Table 1B). Close to the bent zone, the speed ranged from 12.9 to 19.4 cm.s^−1^ between e1 and e3, while it dropped significantly to a range of 5.8 to 3.3 cm.s^−1^ between the farthest electrodes e3 and e6. Triggering and propagation of the electrical signal in response to stem apical bending was found in 100 % of poplars (N= 10).

**Figure 2:**
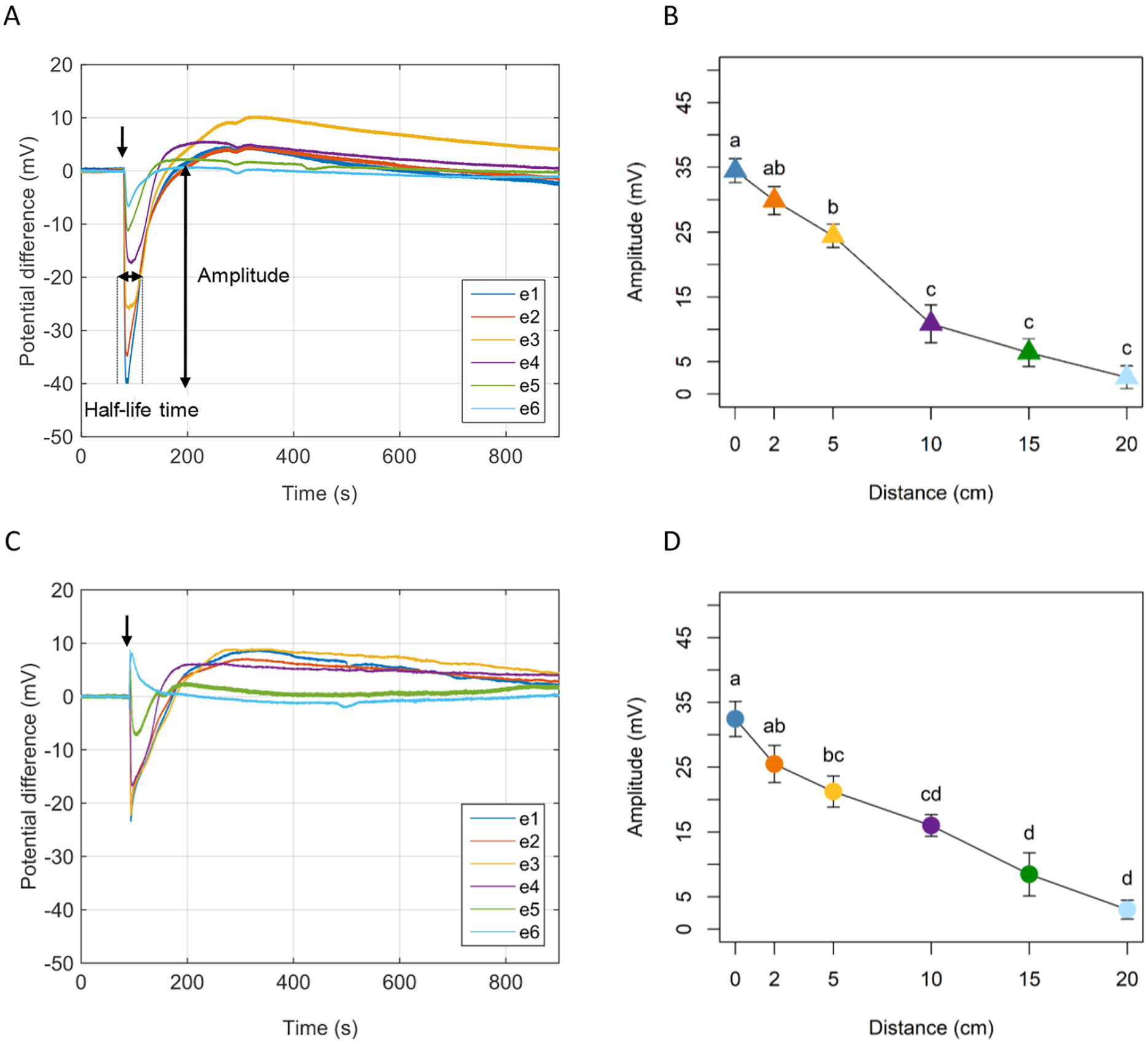
Monitoring of the electrical signal generated after a transient stem bending and recorded at different positions in the stem in *Poplar tremula x alba*. Recording of the electrical signals measured by the six electrodes after a transient apical (A) or basal (C) stem bending applied on the stem (arrow). The half-life time of the signal was defned as the time during which the signal is higher than half the maximal amplitude. Evolution of the amplitude of depolarization wave after apical (B) or basale (D) stem bending depending on the distance from the bent-zone. Values represent the mean (± SE), (n=10). Letters indicate the values that are statistically different (Kruskal and Wallis’s tests and Dunn’s tests p < 0.05).

**Table 1:** Evolution of the half-life time (A) and the mean of the propagation speed (B) of a depolarization wave induced by an apical or basal bending of the stem.

| Electrode number | Half-life time (s) |  |
| --- | --- | --- |
|  | Apical bending | Basal bending |
| e1 | $44.2 \pm 6.7$ | $51 \pm 8.3$ |
| e2 | $32.9 \pm 4.5$ | $43.3 \pm 7.1$ |
| e3 | $34.3 \pm 3.5$ | $42.3 \pm 4.4$ |
| e4 | $21.6 \pm 5$ | $35 \pm 3.6$ |
| e5 | $15.1 \pm 4.9$ | $32.3 \pm 3.9$ |
| e6 | $6.8 \pm 4.4$ | $9.7 \pm 9.7$ |

B
| Interval from the bent-zone (cm) | Average speed (cm.s<sup>-1</sup>) |  |
| --- | --- | --- |
|  | Apical bending | Basal bending |
| [0;2] | $12.98 \pm 2.4$ | $15.12 \pm 5.4$ |
| [2;5] | $19.41 \pm 3.8$ | $16.87 \pm 2.7$ |
| [5;10] | $5.85 \pm 1.7$ | $12.27 \pm 2.7$ |
| [10;15] | $3.31 \pm 1.7$ | $8.43 \pm 1.1$ |
| [15;20] | $3.36 \pm 2.7$ | $1.19 \pm 1.2$ |

**Table 1: Evolution of the half-life time (A) and the mean of the propagation speed (B) of a depolarization wave induced by an apical or basal bending of the stem.** A
| Electrode number | Half-life time (s) |  |
| --- | --- | --- |
|  | Apical bending | Basal bending |
| e1 | 44,2 ± 6,7 | 51 ± 8,3 |
| e2 | 32,9 ± 4,5 | 43,3 ± 7,1 |
| e3 | 34,3 ± 3,5 | 42,3 ± 4,4 |
| e4 | 21,6 ± 5 | 35 ± 3,6 |
| e5 | 15,1 ± 4,9 | 32,3 ± 3,9 |
| e6 | 6,8 ± 4,4 | 9,7 ± 9,7 |

B
| Interval from the bent-zone (cm) | Average speed (cm.s<sup>-1</sup>) |  |
| --- | --- | --- |
|  | Apical bending | Basal bending |
| [0;2] | 12,98 ± 2,4 | 15,12 ± 5,4 |
| [2;5] | 19,41 ± 3,8 | 16,87 ± 2,7 |
| [5;10] | 5,85 ± 1,7 | 12,27 ± 2,7 |
| [10;15] | 3,31 ± 1,7 | 8,43 ± 1,1 |
| [15;20] | 3,36 ± 2,7 | 1,19 ± 1,2 |

A transient and rapid bending of the basal stem segment also generated a depolarization wave (Fig. 2C) propagating acropetally up to 20 cm. The mean amplitude near the bent zone was around 32.4 mV (e1), and dropped to 21.2 mV after 5 cm (e3), to finally reach a mean close to zero (e6 = 3 mV) after 20 cm (Fig. 2D). Following the depolarization phase a slight hyperpolarization was observed on the four measuring electrodes closest to the stimulation. Then, a slow and oscillation-free return to the initial potential occurred. The half-life time of the depolarization wave also decreased with the position of the electrode; from 51 s for e1 to 42.3 s for e3 and 9.7 s for e6 (Table 1A). The average propagation speed of the depolarization wave was evaluated between each electrode. Close to the bent zone, the speed ranged from 15.1 to 16.9 cm.s^−1^ between e1 and e3, while it dropped significantly to a range of 8.4 to 1.2 cm.s^−1^ between the farthest electrodes e3 and e6 (Table 1B). As with the bending of the apical segment, triggering and propagation of the electrical signal in response to stem basal bending was found in 100 % of poplars (N=10).

In order to compare the nature of the electrical signals propagated following different types of stimulation, we therefore tested the response of poplars in the case of a leaf burn. As shown in Fig. 3, a rapid depolarization was observed for each electrode following a leaf burn; starting with those placed closest to the stimulation site. This depolarization wave propagated both acropetally and basipetally in the poplar stem. The return to the initial potential was slow (5 to 45 minutes) and irregular. The amplitude tended to be greater near the stimulation, ranging from 5 to 70 mV. The propagation speed was about 0.02 cm.s^−1^.

**Figure 3:**
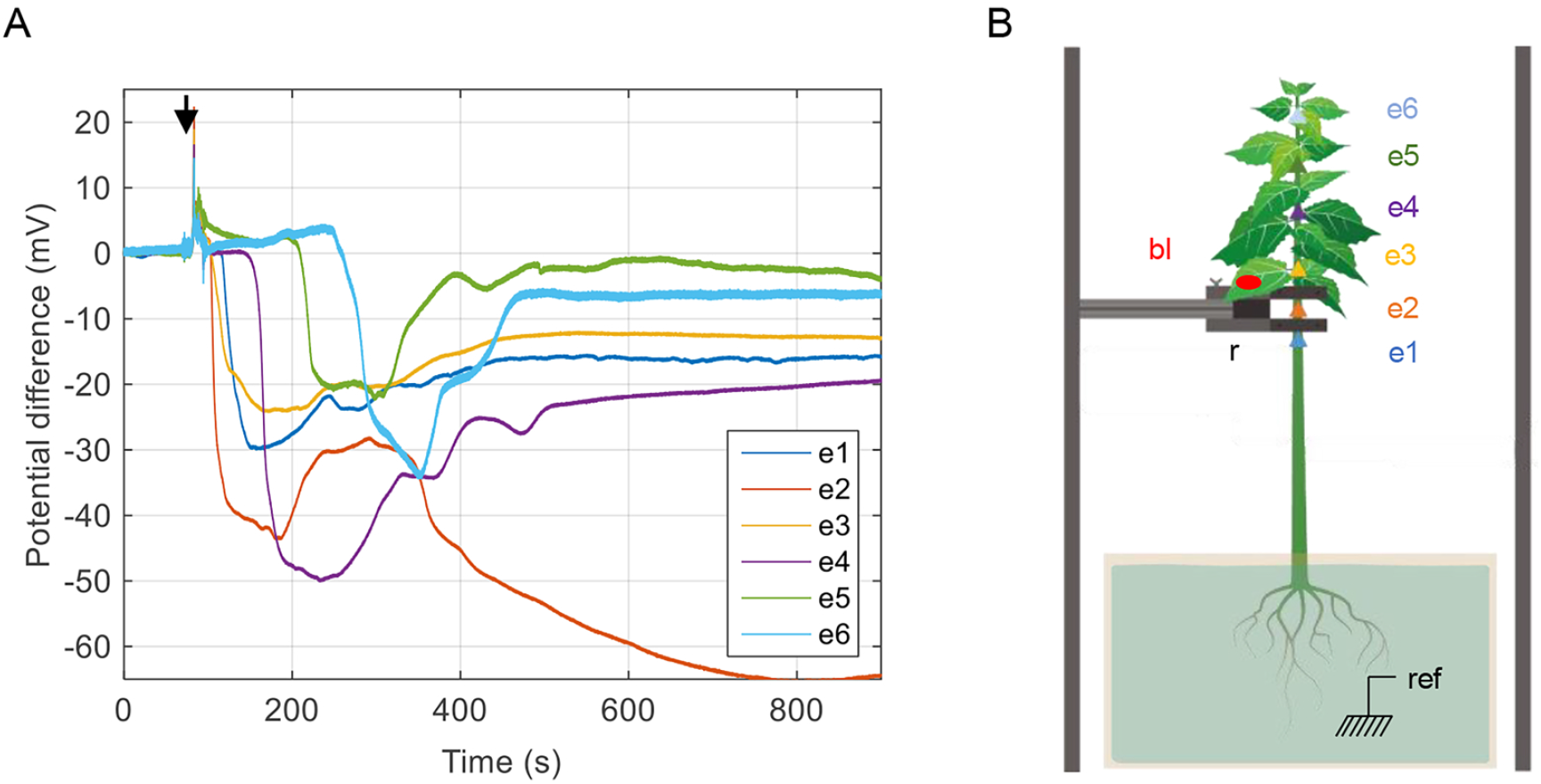
Monitoring of the electrical signal generated after a burnt leaf (bl) in *Poplar tremula x alba.* For each electrode, slow wave consists in a fast depolarization followed by a slow and irregular return to the initial potential. Here the amplitude ranges between 20 to 70 mV and the velocity is around 0.02 cm.s-1(A). The time of the stimulation is indicated by the black arrow. The poplar tree is placed in a Faraday cage; fxed in two points of the stem with clamping rings (r). Stimulation consists of burning the leaf with a match for about 4 s (B). The electrodes are placed along the stem as follows: e1-e2 = 2 cm, e2-e3 = 3 cm, e3-e4 = 5 cm, e4-e5 = 5 cm and e5-e6 = 5 cm.

As complementary experiments, we tried to generate an AP by applying cold water to the stem on eight different poplars individuals (Fig. 4). Whereas three plants did not react, a local depolarization (no propagation) was measured on four of them, with a mean amplitude of 21.3 mV ± 5.5. Only one plant showed a clear depolarization event propagating over a distance of at least 1 cm but less than 3 cm with a velocity of around 0.5 cm.s^−1^ and an amplitude of 30 mV.

**Figure 4:**
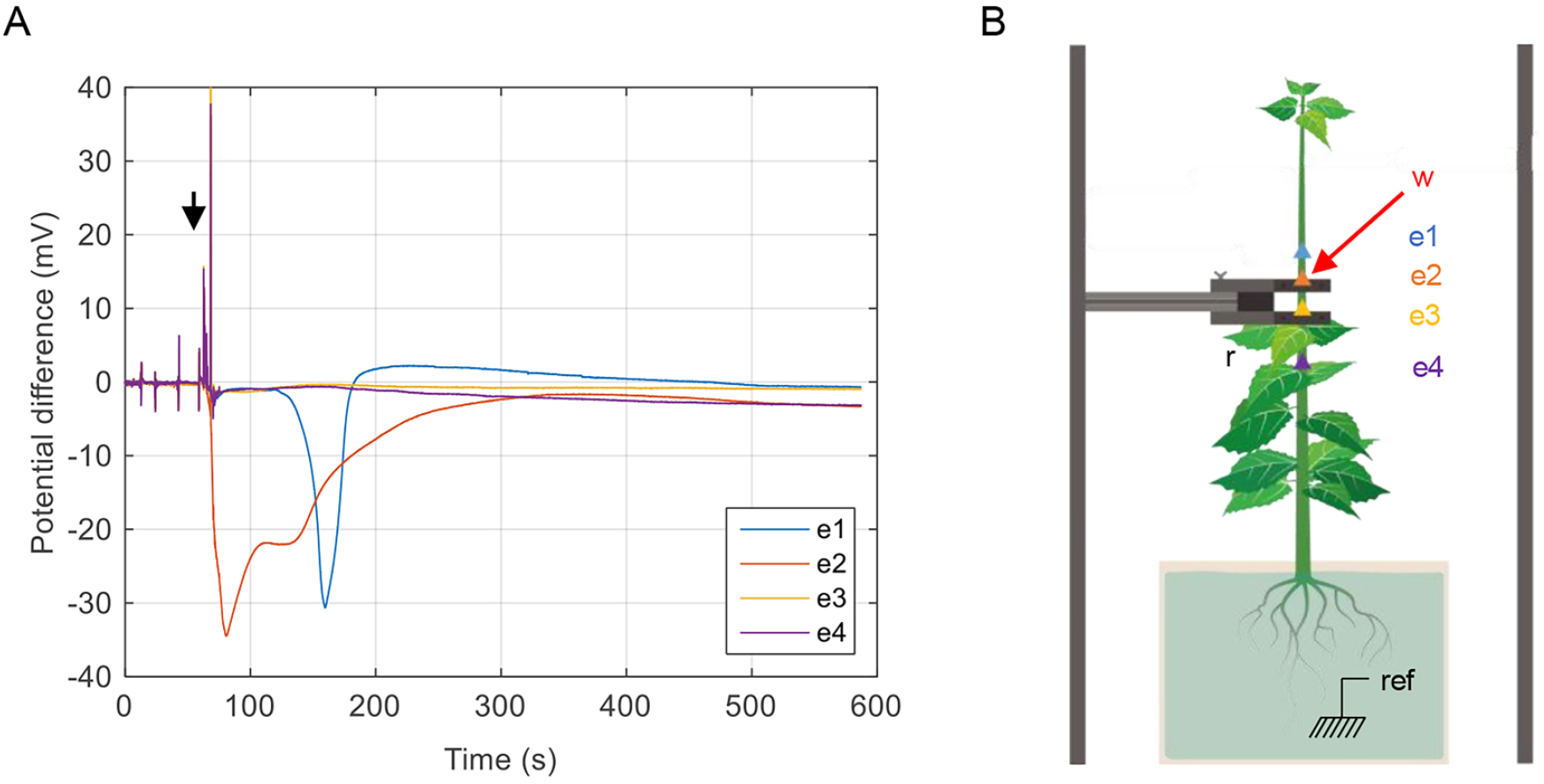
Monitoring of the electrical signal generated after the application of droplet of water at 5°C to the stem (w) in *Poplar tremula x alba.* The time of the stimulation is indicated by the black arrow (A). The poplar tree is placed in a Faraday cage; fxed in two points of the stem with clamping rings (r). The electrodes are placed along the stem as follows: e1-e2 = 2 cm, e2-e3 = 3 cm and e3-e4 = 5 cm (B).

## Discussion

We studied this signaling following three types of stimulations, each triggering electrical signals with different characteristics. In response to a leaf burn, a depolarization typical of a SW was generated in the poplar trees (Fig. 3). The characteristics of this SW are identical than those described in the literature for herbaceous species: a fast depolarization that tends to decrease with distance followed by a slow and irregular return to the initial potential and a propagation velocity of the order of 0.01cm.s^−1^ (Sambeek and Pickard, 1976; Roblin and Bonnemain, 1985; Julien et al., 1991). On the one hand, the recording of the SW in response to wounding allows validating our experimental setup used for measuring the propagation of electrical signals in poplar. On the other hand, the comparison of the two electrical responses clearly suggests that the one caused by the bending of the stem is not a slow wave.

The shape of the electrical response induced by stem bending, a rapid depolarization followed by a repolarization at the initial potential, shows similarities with AP as described by Pickard, 1973. The maximum amplitude reached near the bent zone was around 34 mV; a low value compared to AP measured in various plant species. The position of the six electrodes along the stem allowed following the electrical signal over a distance of 20 cm. These measurements enlightened a significant attenuation of the amplitude and velocity with the increasing distance from the stimulation point (Figs. 2B; 2D and Table 1B). On the contrary, the literature reports that the plant AP propagates at constant rate and without decrement (Sukhov et al., 2018). Although very few studies have focused on studying it over a distance greater than 10 cm, this was observed by Zawadzki et al. (1991) where an AP electrically generated on a stem of *Helianthus annuus* propagates over at least 15 cm of stem at constant rate and without decrement, later confirmed by Stankovic et al. (1998). This decrement is a first major difference that distinguishes our signal from a classical plant AP.

Despite its action potential-like shape, the electrical response recorded following stem bending does not match with the definition of AP: it differs by the slow decreasing of the amplitude during the propagation of the signal. Morever, it differs from the cold induced putative action potential described in *Populus trichocarpa* by a speed 30 times higher (Lautner et al., 2005). Furthermore, the amplitude of the AP observed by Lautner et al., (2005) did not seem to show a decrement, although it is not specified if the distance between the two electrodes was higher or less than 10 cm. To take into account this typical characteristic of bending-induced electrical response, we propose to descriptively call this electrical signal as a ‘gradual’ potential.

We propose two hypotheses to explain the decrement of the ‘gradual’ potential. The first hypothesis assumes an electrotonic propagation of the ‘attenuated’ potential. In this case, the decrement can be explained by the poor performance of the excitable and connected cells (especially the phloem) as a propagation medium for an electrical signal. Indeed, the electrical resistance caused by the shrinkage of the membrane at the plasmodesmata, and the capacitance of the non-myelinated membrane (companion cells) cause the dissipation of the electrical signal (Hedrich et al., 2016). The possible re-amplification of the signal along the phloem by voltage-dependent channels would then not be sufficient to compensate for this dissipation. Indeed, no equivalent to Ranvier’s nodes has been identified in plants to date. The transmission of the excited state to neighboring cells would tend to decrease, which could account for the drop in propagation speed. The second hypothesis suggests that the ‘gradual’ potential observed following a transient bending of the stem could result from the opening of mechanosensitive channels following the passage of a hydraulic pressure wave in the vascular tissues, itself induced by the bending (Lopez *et* al., 2014, Louf et al., 2017). This hydraulic pulse is itself attenuated over the distance because of lateral leaks and the mechanical flexibility of the pipes, which absorb some of the initial energy.

## Conclusion

Transient bending of poplar stems generates a rapid depolarization wave that to propagates rapidly up to 20 cm from the stimulation zone with decrement. The characteristics of this depolarization wave do not fit with the definitions neither of a slow wave nor of the well-known action potential. Also, we propose to call this signal ‘gradual’ potential. Its propagation mechanism remains unknown. That is why current research is now focusing on the hypothesis of hydro-electric coupling.

## Materials and methods

### Plant material and culture conditions

Young poplars (*Populus tremula×alba*, clone INRA 717-1B4) were obtained by *in vitro* micropropagation (Leple et al., 1992). Once they reached a height of about 4 cm, they were gradually acclimatized on a hydroponic solution (Martin et al., 2009) through decreasing relative humidity. Trees were then placed in a growth chamber (16 h/8 h light/dark cycle at 40 μmol.m^−2^.s^−1^ and 22 °C/18 °C with air relative humidity of 60%). Four months after micropropagation, the poplars were used for experiments: at this stage, stems were about 77.8 cm (±1.5 SE) tall with an average diameter of 5.8 mm (±0.3 SE).

Prior to electrophysiology experiments plants were moved into a Faraday cage in ambient laboratory (16 h/8h light/dark cycle at 20 μmol.m^−2^.s^−1^ and 22 °C/20 °C). Plants were set vertically and fixed in two points of the stem with clamping rings (Fig. 1). Foam was rolled around each part of the stem before tightening the clamping rings to avoid stem wounds and to allow possible stem diameter variations. The root system was plunged in a 20 L vat filled with hydroponic solution.

### Bending treatments

The leaves of the bent segment were removed with a razor blade to avoid uncontrolled mechanical stimuli. The speed and the magnitude of the bending stimulation were controlled by a motorized arm (hybrid stepping motors 17PM-H311-P1, Minebea.Co) that pushed the stem against a plastic template; which had a constant radius of curvature. The speed of the motor was fixed in order to manage a flexion time of 2 s (round trip) using interface software MegunoLink. The strain magnitude on the stem periphery was controlled by the radius of curvature of the plastic template. This method allowed applying the same strain level along 12 cm of the bent segment. The setup was adjusted in order to apply a maximal peripheral longitudinal strain of the bark of around 2% (Moulia et al., 2015). This value is high enough to generate significant thigmomorphogenetic responses (Niez et al., 2019).

Two types of bending treatments were applied. For transient apical bending, the template was placed 35 cm below the apex (Fig. 1A). For transient basal bending, the template was placed 25 cm above the roots (Fig. 1B).

### Burning treatment

The burn was carried out at a leaflet along the midrib with a match. The flame was held 1 cm below the leaflet for 4 to 6 seconds.

### Cold treatment

The cold treatment was applied by placing a drop of cold water (5°C) with a pipet tip near the upper clamping ring.

### Monitoring of extracellular electrical signals

In order to minimize artefactual interferences, one single poplar was placed in a Faraday cage. All electronic materials were located outside the cage. Six measuring electrodes (tinned copper wire (359-835), 0.25 mm diameter, RS Components) were inserted into the stem, passing through all the tissue to the pith, and measured electrical potential simultaneously near the bending area and up to 20 cm from it. After the installation was completed, the plant was left undisturbed for stabilization during 24 to 48 hours.

The reference electrode (RC3 model, World Precision Instruments) was made of an Ag/AgCl wire immersed in the nutrient solution of the plants. The electrical potential recorded is the difference between a measuring electrode and the reference electrode with respect to ground. The first measuring electrode e1 was inserted 2 cm below (apical bending) or above (basal bending) the stem-to-template contact (Fig. 1). The respective distances of each electrode from e1 were 2 cm (e2), 5 cm (e3), 10 cm (e4), 15 cm (e5), and 20 cm (e6). The six measuring electrodes and the reference electrode were connected to a cDAQ-9171 (National Instrument) electronic card. The electronic card was used as an impedance amplifier (10 GΩ) and A/D converter. DAQExpress 1.0.1 software (National Instrument) monitored the potential difference with a sampling rate of 200 Hz. The graphs and analyzes were built using MatLab® software.

The analysis of the recordings provided several parameters of the signal. The amplitude was defined as the maximum difference value compared to the baseline before bending. The half-life time of the signal was defined as the duration at half amplitude (Fig. 2A). The average propagation speed of the signal was calculated between each electrode as the ratio of the distance between two consecutive electrodes to the delay between the detection of a change in potential difference.

### Statistical analysis

All measured and computed data were statistically analysed using R software. Kruskal and Wallis’s tests and Dunn’s tests were performed to compare results in terms of amplitude, duration and speed of the signal (p < 0.05).

## Acknowledgments

The authors thank Brigitte Girard, Romain Souchal, Stephane Ploquin, Amélie Coston and Juliette Tinturier for their technical support. And thank Rémi Cadet, Jean-Marie Frachisse, Yoël Forterre and Bertrand Coste for the fruitful discussions.

